# Electrical stimulation enhances carbon tetrachloride removal in a simulated wetland–groundwater interaction zone

**DOI:** 10.64898/2026.11.26.768792

**Authors:** Xin Chen, Jiahao Zhang, Fengyuan Zhuang, Siming Chen

## Abstract

Carbon tetrachloride (CCl_4_) contamination presents persistent challenges for groundwater remediation. This study evaluated electrical stimulation and plant–microorganism interactions in sediment columns simulating a wetland–groundwater interaction zone. Five treatments were compared at initial CCl_4_ concentrations of 2, 5 and 10 mg/L, with a cell voltage of 0.6 V applied to electrically stimulated reactors. At 2 mg/L, non-autoclaved sediment treatments achieved 79.5–82.9% removal within 48 h, compared with 47.3–61.1% in autoclaved-sediment treatments. Nearly complete removal occurred in both electrically stimulated treatments during the first cycle at 10 mg/L, and high removal persisted across subsequent cycles. Chloride concentrations increased alongside CCl_4_ removal, while bulk organic carbon declined. The pH remained predominantly between 7.1 and 8.1. Sequencing revealed treatment-associated community differences. The *Pseudomonas* increased from 36.1% in the inoculum to 55.8% in the planted, electrically stimulated treatment at 2 mg/L. Electrode-associated *Methylobacillus* and *Methylophilus* characterized unplanted electrically stimulated reactors, whereas *Methylotenera* characterized planted counterparts. These findings demonstrate sustained removal and identify microbial populations potentially associated with treatment performance. The results support further evaluation of electrically assisted planted systems for groundwater remediation at shallow interfaces accessible to roots.

## 1. Introduction

Groundwater accounts for approximately 99% of the Earth’s liquid freshwater resources and supplies approximately 50% of global domestic water withdrawals and 25% of agricultural irrigation withdrawals, making its protection essential for water security and groundwater-dependent ecosystems (UNESCO World Water Assessment Programme, 2022). Carbon tetrachloride (CCl_4_), a volatile chlorinated compound historically used in industrial processes and grain fumigation, remains a concern at legacy contamination sites because of its toxicity and persistent groundwater contamination. Historical investigations reported groundwater concentrations of approximately 940 μg/L at Waverly, Nebraska, and 910 μg/L in a well at Hilton, Kansas (LaFreniere, 2008; U.S. Environmental Protection Agency). These concentrations greatly exceed the WHO drinking-water guideline value of 4 μg/L, the U.S. maximum contaminant level of 5 μg/L, and the Chinese drinking-water limit of 2 μg/L. Developing effective treatment approaches for contaminated groundwater, including at interfaces with wetland sediments, is therefore an important remediation objective.

A variety of physical, chemical, and biological remediation technologies have been developed for CCl_4_-contaminated groundwater, including zero-valent iron permeable reactive barriers, nanoscale zero-valent iron injection, in situ chemical reduction, and enhanced reductive dichlorination (Lawrinenko et al., 2023; Ling et al., 2024; Liu et al., 2025). Reductive transformation is an important route for CCl_4_ removal, but its products depend on the reaction conditions. For example, sulfide-containing catalytic systems can generate chloroform, carbon disulfide, bicarbonate and chloride (Ding et al., 2022). However, treatment performance can be constrained by both geochemical and biological factors. In iron-based barriers, surface passivation and secondary mineral accumulation can reduce reactivity and alter hydraulic properties (Lawrinenko et al., 2023). Biological reduction of CCl_4_ is also highly dependent on the availability of effective electron donors, suitable redox conditions, and microorganisms capable of the relevant transformations. For instance, organic electron-donor supplementation promoted CCl_4_ removal in groundwater microcosms (Liu et al., 2025). Removal of the parent compound nevertheless needs to be distinguished from complete dechlorination and mineralization, because chlorinated intermediates can require additional microbial transformations (Fernández-Verdejo et al., 2022b; Zhu et al., 2022).

Bioelectrochemical systems (BESs) provide an approach to managing electron availability during groundwater remediation. Depending on electrode potential and microbial physiology, cathodic electrons may support dechlorination through direct or mediated electron transfer, or through hydrogen generated at the cathode (Wang et al., 2025; Zhu et al., 2022). Experimental evidence illustrates the importance of complementary microbial functions: in a bioelectrochemical co-culture, cathodically generated hydrogen supported chloroform reduction to dichloromethane by *Dehalobacter*, followed by dichloromethane fermentation to acetate by *Dehalobacterium* (Fernández-Verdejo et al., 2022b). Permeable bioelectrochemical reactors have also demonstrated the feasibility of integrating electrochemical assistance into groundwater treatment configurations (Sassetto et al., 2024). These findings provide a rationale for electrical stimulation, while emphasizing that treatment outcomes depend on the transformation pathways and microbial functions present in each system.

Plant roots provide a further means of modifying the local conditions that influence contaminant transformation. In reeds, internal oxygen transport and radial oxygen loss can generate spatially and temporally variable oxygen distributions in otherwise oxygen-depleted sediments (Li et al., 2021). Root exudates and root surfaces can also supply organic substrates and habitats for microbial growth, contributing to the treatment functions of constructed wetlands (Wu et al., 2023). The relevance of planted systems to chlorinated groundwater contamination is illustrated by phytoremediation research on trichloroethylene and a full-scale reedbed treating groundwater contaminated with 1,2-dichloroethane (Cheng et al., 2024; Rahim et al., 2026). Together, these studies motivate testing whether electrical stimulation can improve CCl_4_ removal in planted wetland sediments. A potential mechanism is the coexistence of reducing conditions that support dechlorination and root-influenced microenvironments that support subsequent transformations. However, such spatial coupling remains a hypothesis for CCl_4_ treatment and requires direct evidence of local redox conditions and transformation products.

Therefore, column reactors were constructed in this study to investigate the effects of electrical stimulation on CCl_4_ removal in a simulated wetland–groundwater interaction zone containing reeds and indigenous sediment microorganisms. Five treatment groups were compared at initial CCl_4_ concentrations of 2, 5 and 10 mg/L to evaluate removal performance and treatment-associated differences. CCl_4_ concentrations, chloride release, organic carbon, pH and oxidation–reduction potential were monitored to assess contaminant removal, indicators of dechlorination and changes in bulk water chemistry. The 16S rRNA gene sequencing characterized microbial community responses in bulk sediment. These measurements were used to evaluate treatment performance and develop hypotheses about the processes contributing to CCl_4_ removal. This study provides an experimental basis for assessing the potential of electrically assisted plant–microorganism systems for groundwater remediation.

## 2. Materials and Methods

### 2.1. Chemicals

All reagents were of analytical grade or higher and were purchased from Sinopharm Chemical Reagent Co., Ltd. or Shanghai Macklin Biochemical Co., Ltd. Carbon tetrachloride (HPLC grade, purity ≥99.9%) was used as the target contaminant. HPLC-grade methanol was used to prepare a CCl_4_ stock solution at 1,000 mg/L. All aqueous solutions were prepared using ultrapure water. Synthetic groundwater was prepared using a previously reported formulation (Chen et al., 2023). Its composition, including calcium, magnesium, sodium, potassium, chloride, bicarbonate and phosphate, is provided in Table 1. The pH was adjusted to 7.2 ± 0.1 using 0.1 mol/L HCl or NaOH. Before use, the synthetic groundwater was deoxygenated by sparing N_2_ for 20 minutes and amended with the CCl_4_ stock solution to achieve target concentrations of 2.0, 5.0 and 10 mg/L. No other organic electron donors were deliberately added.

**Table 1.** Chemical composition of synthetic groundwater.

| Chemicals | Concentration (g/L) |
|---|---|
| CaCl<sub>2</sub> | 0.2464 |
| MgCl<sub>2</sub>·H<sub>2</sub>O | 1.0572 |
| NaCl | 0.4459 |
| KCl | 0.0283 |
| NaHCO<sub>3</sub> | 0.8082 |
| KH<sub>2</sub>PO<sub>4</sub> | 0.0299 |

### 2.2. Inoculant

Wetland sediment was collected from a site with a history of chlorinated-hydrocarbon contamination in Jiangsu, China. Previous site investigations reported sediment chlorinated-hydrocarbon concentrations ranging from 0 to 6809.1 mg/kg. Sediment was collected using sterile tools, transferred to sterile polyethylene bags and transported to the laboratory in an insulated container with ice packs. Visible plant debris, stones and other coarse materials were removed. The sediment was homogenized, passed through a sterile 2.0-mm nylon sieve and stored at 4°C until reactor preparation. A portion of the sediment was autoclaved at 121°C for 30 min twice. This autoclaved sediment was used to prepare the control treatments.

### 2.3. Wetland-groundwater interaction zone column reactor construction

A customized vertical column reactor was used to construct wetland-groundwater interactions zone. The reactor body was made of transparent polymethyl methacrylate, with an inner diameter of 100 mm, an outer diameter of 110 mm, and a total height of 220 mm. Groundwater inlet and outlet channels were installed to simulate continuous groundwater flow. The internal column was vertically divided into three functional zones, including aquifer, wetland-groundwater interaction zone, and wetland sediment zone (Fig. 1). A water layer of approximately 20 mm was maintained above the sediment to simulate natural inundated wetland conditions. During experimental operation, the outer surface of each reactor was wrapped with two layers of aluminum foil to prevent algal growth induced by light exposure and to minimize potential effects of light on CCl_4_ volatilization. The foil was removed only briefly during observation and maintenance. Synthetic groundwater containing CCl_4_ was introduced from the bottom of the column, and the influent flow rate was controlled using a peristaltic pump. The reactors were operated continuously. All reactors were maintained at 25 ± 2 °C under dark conditions. The corresponding influent flow rate was approximately 65 mL/h.

**Figure 1.**
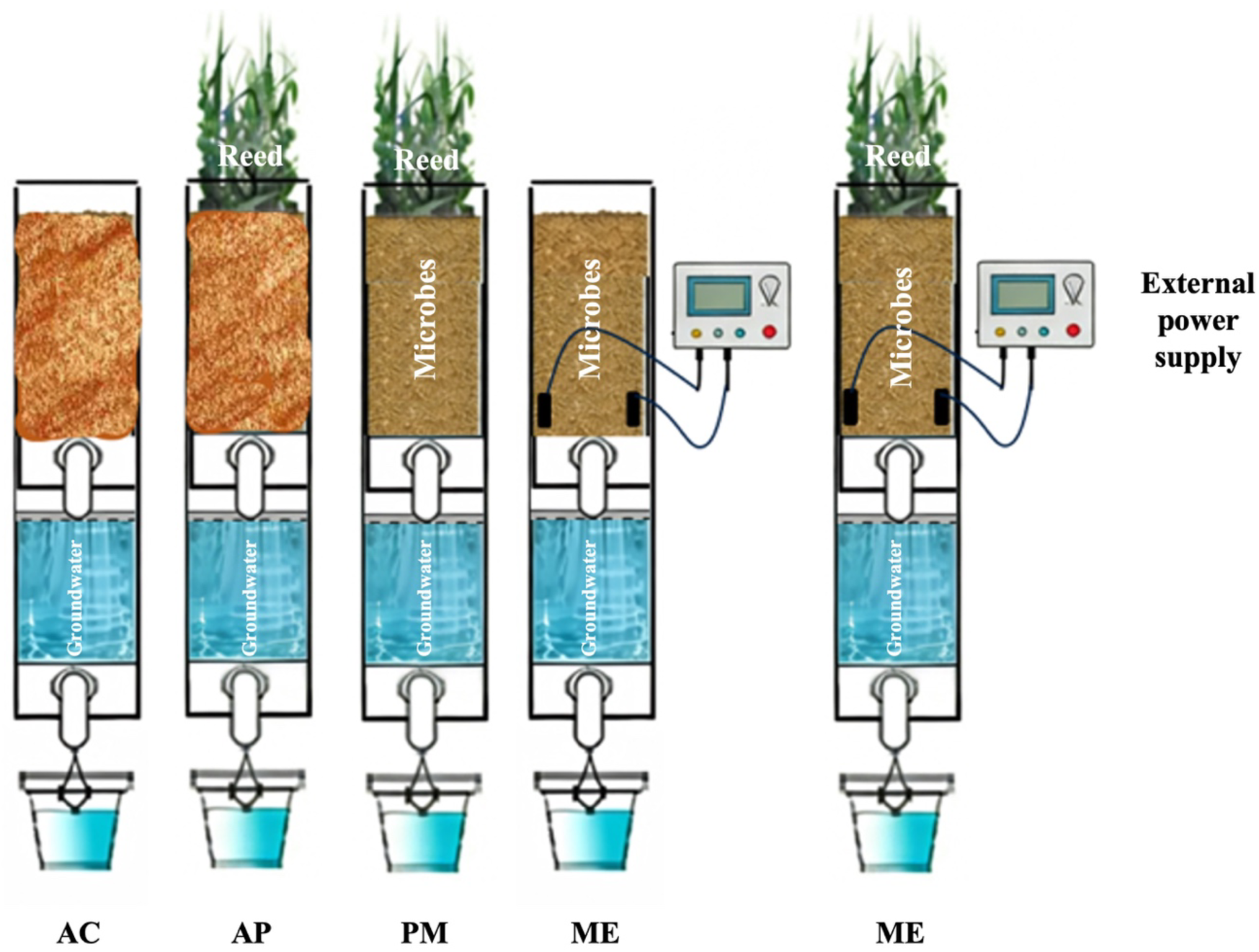
Schematic of the experimental column reactors.

Graphite plates were used as the anode and cathode in this study due to their good electrical conductivity, chemical stability, and biocompatibility.(Singh et al., 2022) Each graphite electrode measured 40 mm × 20 mm × 5 mm, with an effective surface area of approximately 10 cm². The electrodes were vertically and parallelly positioned in the wetland–groundwater interaction zone (150–170 mm above the column bottom), with an electrode spacing of 10 mm. The upper ends of the electrodes were connected to corrosion-resistant titanium wires using conductive epoxy resin and sealed with waterproof silicone to prevent water leakage and exposure of the metallic connection areas. A DC power supply maintained a constant cell voltage of 0.6 V between the anode and cathode in the electrically stimulated treatments (Chen et al., 2023; Marshall et al., 2012). Reed was used in this study because of its strong environmental adaptability, and tolerance to chlorinated organic contaminants. Before the experiment, the root surfaces were gently rinsed with deionized water to remove attached sediment particles and residual disinfectant. The treated reed seedlings were acclimated under controlled environmental conditions for 3–5 d, during which synthetic groundwater was used to maintain root-zone adaptation. The initial fresh weight of selected plants ranged from 30–45 g. The number of seedlings planted in each column was four. Five experimental treatment groups were established (Table 2). AC provided an autoclaved-sediment control, whereas AP assessed removal in planted columns containing autoclaved sediment. PM, ME and PME were used to compare removal in plant–microorganism, electrically stimulated microorganism, and combined plant–microorganism–electrical systems, respectively.

**Table 2.** Experimental setup.

| Experimental group | Treatment | Electric field | Plant |
|---|---|---|---|
| AC | Autoclaved control | - | - |
| AP | Plants | - | + |
| PM | Plants, Microbes | - | + |
| ME | Microbes, Electric field | + | + |
| PME | Plants, Microbes, Electric field | + | + |

### 2.4. Chemical Analysis

During reactor operation, water samples were collected from sampling ports at different heights using gas-tight syringes to minimize losses of CCl_4_ and its volatile transformation intermediates during sampling. Before each sampling event, approximately 10 mL of residual water in the sampling tubing was slowly withdrawn and discarded to eliminate the influence of sampling-line dead volume. Formal analytical samples were then collected. The samples were immediately transferred into pre-cleaned glass vials specifically designed for volatile organic compounds. The vials were sealed with polytetrafluoroethylene (PTFE)-lined septa, and headspace bubbles were avoided to minimize volatilization losses. All samples were stored at 4°C in the dark and analyzed within 24 h. Experimental blanks, replicate samples, and calibration standards were included throughout the sampling and analytical procedures to ensure analytical accuracy and reproducibility.

CCl_4_ were determined using headspace gas chromatography (Agilent 6890-5973). The coefficient of determination of each calibration curve was required to exceed 0.995. Standard solutions were freshly prepared or stored for only a short period at 4 °C in the dark. Before each analytical run, the instrument was checked using a blank to confirm the absence of obvious residual contamination before sample analysis. The pH and oxidation-reduction potential (ORP) were measured using a multiparameter water-quality analyzer (DZB-712, Leici). Chloride ion (Cl⁻) concentrations were determined by ion chromatography. Total organic carbon (TOC) was measured using a TOC analyzer (Multi N/C 2100, Analytik Jena). Water samples were filtered through 0.45-μm membrane filters prior to analysis to remove suspended particulate matter.

### 2.5. Microbial Analysis

At the end of the experiment, bulk sediment samples were collected from the different treatment groups for microbial community structure analysis. All samples were collected using aseptic procedures to avoid cross-contamination. After collection, the samples were immediately placed in sterile centrifuge tubes, rapidly transferred to liquid nitrogen, and subsequently stored at −80°C until DNA extraction. Total microbial DNA was extracted from sediment samples and analyzed as described before (Li et al., 2026). DNA concentration and purity were measured using a NanoDrop 2000 spectrophotometer, and DNA integrity was assessed by 1% agarose gel electrophoresis. The V3–V4 hypervariable region of the 16S rRNA gene was selected for amplification using the universal primers 341F (5′-CCTACGGGNGGCWGCAG-3′) and 806R (5′-GGACTACHVGGGTWTCTAAT-3′).(Walters et al., 2016). Raw sequences obtained by high-throughput sequencing were first demultiplexed according to sample barcode information, after which barcode and primer sequences were removed. Quality control was then performed. The resulting high-quality sequences were then imported into the QIIME 2 platform for subsequent analysis.(Bolyen et al., 2019; Caporaso et al., 2010) The DADA2 algorithm was used to denoise the high-quality sequences, including sequence quality filtering, sequencing-error correction, chimera removal, and generation of amplicon sequence variants (ASVs).(Callahan et al., 2016).

### 2.6. Statistical Analysis

CCl_4_ removal efficiency, Cl^-^ release, pH and ORP were statistically compared among different treatment groups. Microbial community data were analyzed using QIIME 2.

## 3. Results

### 3.1. CCl₄ removal and changes in chloride and organic carbon

CCl₄ concentrations decreased in all treatments over the experimental period. At an initial concentration of 2 mg/L, removal efficiencies in PM, ME and PME were significantly higher than those in AC and AP (p < 0.05; Fig. 2a). Removal was particularly rapid during the first 24 h, after which only 32.2–36.7% of the initial CCl_4_ remained in PM, ME and PME. By 48 h, removal efficiencies in these treatments reached 79.5–82.9%, compared with 47.3% in AC and 61.1% in AP. The higher removal in treatments containing non-autoclaved sediment was consistent with a microbial contribution to CCl_4_ removal. However, the substantial concentration decreases in the control treatments indicate that aqueous CCl_4_ loss cannot be attributed exclusively to biodegradation.

**Figure 2.**
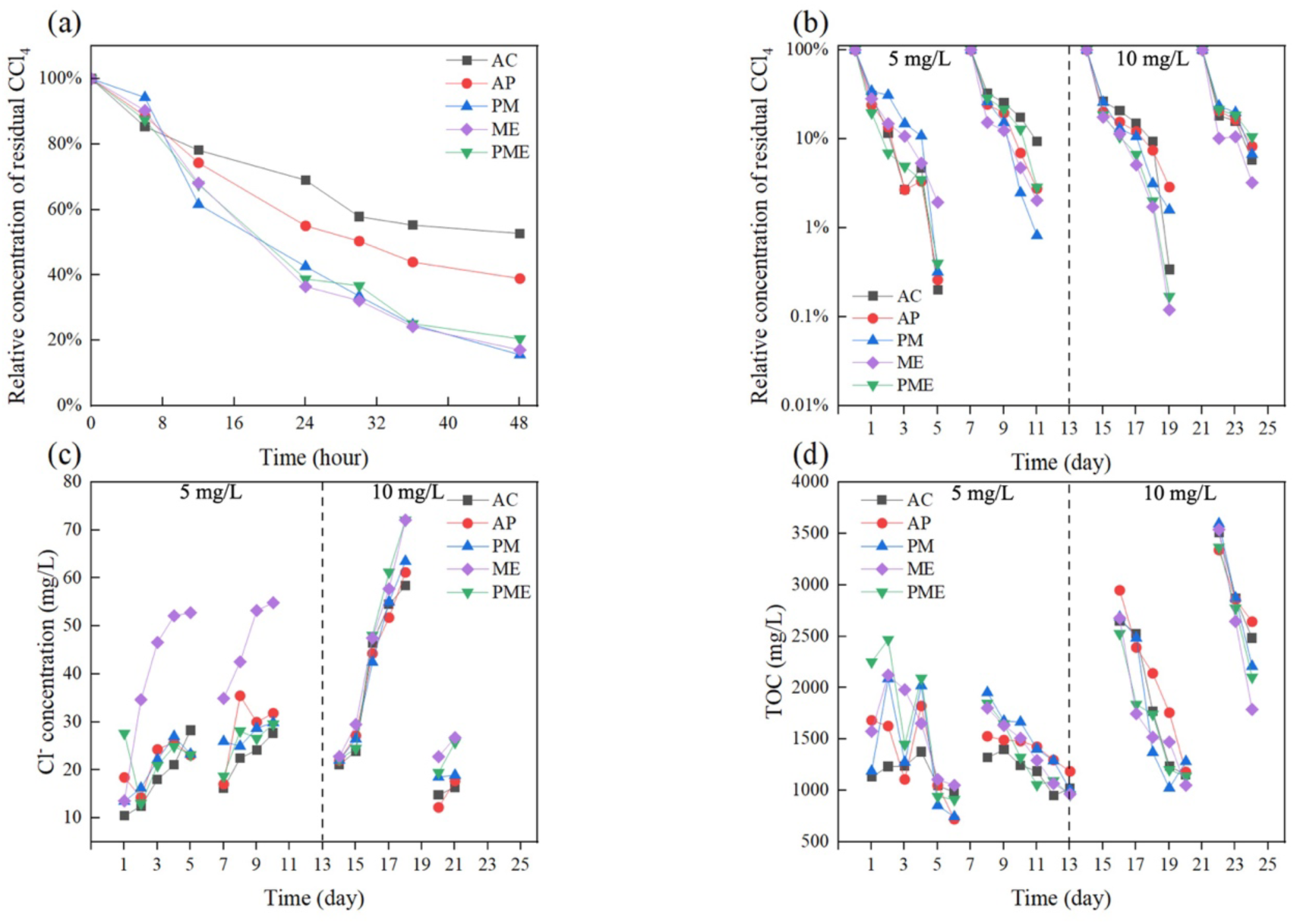
CCl_4_ removal, chloride release and total organic carbon dynamics in the column reactors. Relative concentration of residual CCl_4_ at initial concentrations of (a) 2 mg/L and (b) 5 and 10 mg/L. Temporal changes in (c) chloride ion and (d) TOC concentrations during operation at initial CCl_4_ concentrations of 5 and 10 mg/L.

The reactors were subsequently evaluated over repeated cycles at higher initial CCl_4_ concentrations of 5 and 10 mg/L (Fig. 2b). Nearly complete removal was observed in ME and PME during the first cycle at 10 mg/L. Both electrically stimulated treatments maintained high removal efficiencies over subsequent cycles, demonstrating sustained treatment performance under the tested conditions. Chloride concentrations increased as CCl_4_ concentrations declined (Fig. 2c). The increases were particularly pronounced in ME and PME, where chloride concentrations reached 71.2 and 63.5 mg/L, respectively. Lower chloride concentrations were observed in the other treatments, particularly those containing autoclaved sediment. This pattern is consistent with a contribution from dechlorination, although its extent cannot be quantified without correcting for background chloride and accounting for cumulative inputs and outputs.

TOC concentrations also decreased during reactor operation (Fig. 2d). Averaged across all five treatments, TOC decreased from 1628.9 ± 353.6 to 955.5 ± 280.8 mg/L at an initial CCl_4_ concentration of 5 mg/L, and from 3084.4 ± 426.0 to 1391.1 ± 304.7 mg/L at 10 mg/L. ME and PME consistently exhibited lower TOC concentrations than the other treatments over the four cycles. These observations show that CCl_4_ removal was accompanied by chloride accumulation and bulk organic-carbon removal. The TOC changes cannot, however, establish CCl_4_-specific mineralization because the measured carbon pool also included other sources, including methanol introduced with the CCl_4_ stock solution.

### 3.2. pH and redox conditions during reactor operation

The pH remained predominantly between 7.1 and 8.1 throughout reactor operation (Fig. 3a). Thus, CCl_4_ removal occurred under near-neutral to mildly alkaline conditions, without large excursions in bulk pH during electrical stimulation at a cell voltage of 0.6 V. Bulk ORP values ranged from −75 to −20 mV, with slight increases observed at higher CCl_4_ concentrations (Fig. 3b). During the second cycle at 5 mg/L, ORP decreased in ME and PME to −57 and −71 mV, respectively. At 10 mg/L, AP exhibited a slightly higher ORP than AC. This difference may reflect plant-associated modification of local redox conditions. Previous measurements in Phragmites australis have demonstrated spatially variable rhizosphere oxygen release that depends on root development and environmental conditions (Li et al., 2021).

**Figure 3.**
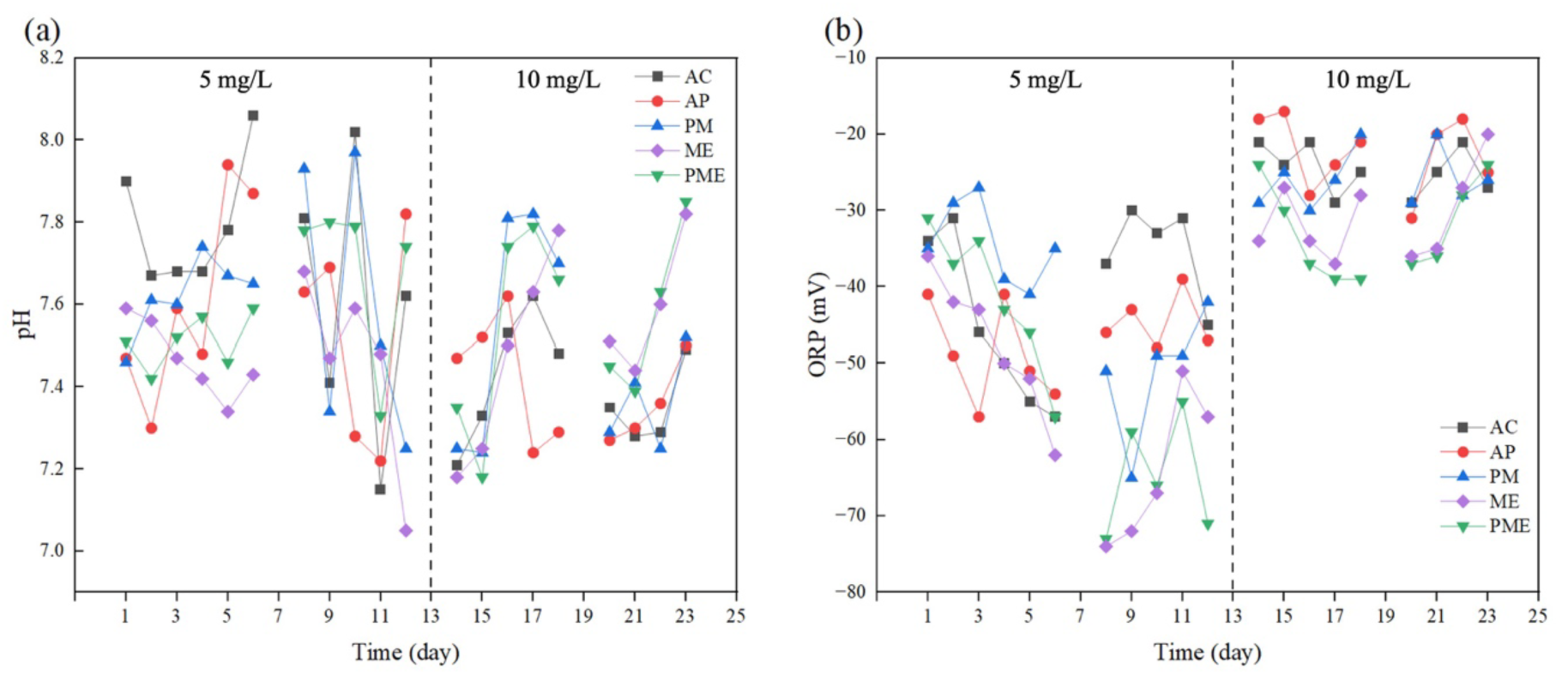
Physicochemical conditions in the column reactors. (a) pH; (b) ORP.

### 3.3. Microbial community structure in bulk sediment and on electrodes

16S rRNA gene sequencing revealed differences in microbial community composition among treatments and CCl_4_ concentration stages. *Pseudomonas* was the dominant genus in the inoculum, accounting for 36.1% of the community, and its relative abundance increased to 55.8% in PME at the end of the experiment with an initial CCl_4_ concentration of 2 mg/L (Fig. 4a). At 5 mg/L, *Methylophaga* and *Halomonas* were prominent in PME, representing 38.3% and 9.5% of the community, respectively (Fig. 4b). Fisher’s exact tests indicated higher representation of *Pseudomonas* in ME and PME than in PM at 2 mg/L (*p* < 0.05; Fig. 4c,d). Conversely, *Methylotenera* was relatively more abundant in PM than in ME and PME at 5 mg/L (*p* < 0.05; Fig. 4e,f). Analysis of electrode-associated communities further revealed distinct taxonomic patterns. *Methylobacillus* and *Methylophilus* were characteristic taxa on ME electrodes (Fig. 5a,b), whereas *Methylotenera* was distinctively distributed in the PME electrode (Fig. 5c,d). These observations suggest that plant presence was associated with differences in the composition of electrode-associated communities, extending the community patterns observed in bulk sediment. Electrode association alone, however, does not demonstrate direct electron exchange with the electrode. The observed community composition provides a basis for proposing potential microbial contributions to CCl₄ removal (Fig. 6). For example, the strain historically designated *Pseudomonas stutzeri* KC transforms CCl₄ through reactions associated with pyridine-2,6-bis(thiocarboxylic acid) (PDTC), and its PDTC biosynthesis genes occur on an integrative and conjugative element (Sewell et al., 2024). This strain-specific precedent supports investigating a potential role for the detected *Pseudomonas*-affiliated populations in CCl_4_ transformation.

**Figure 4.**
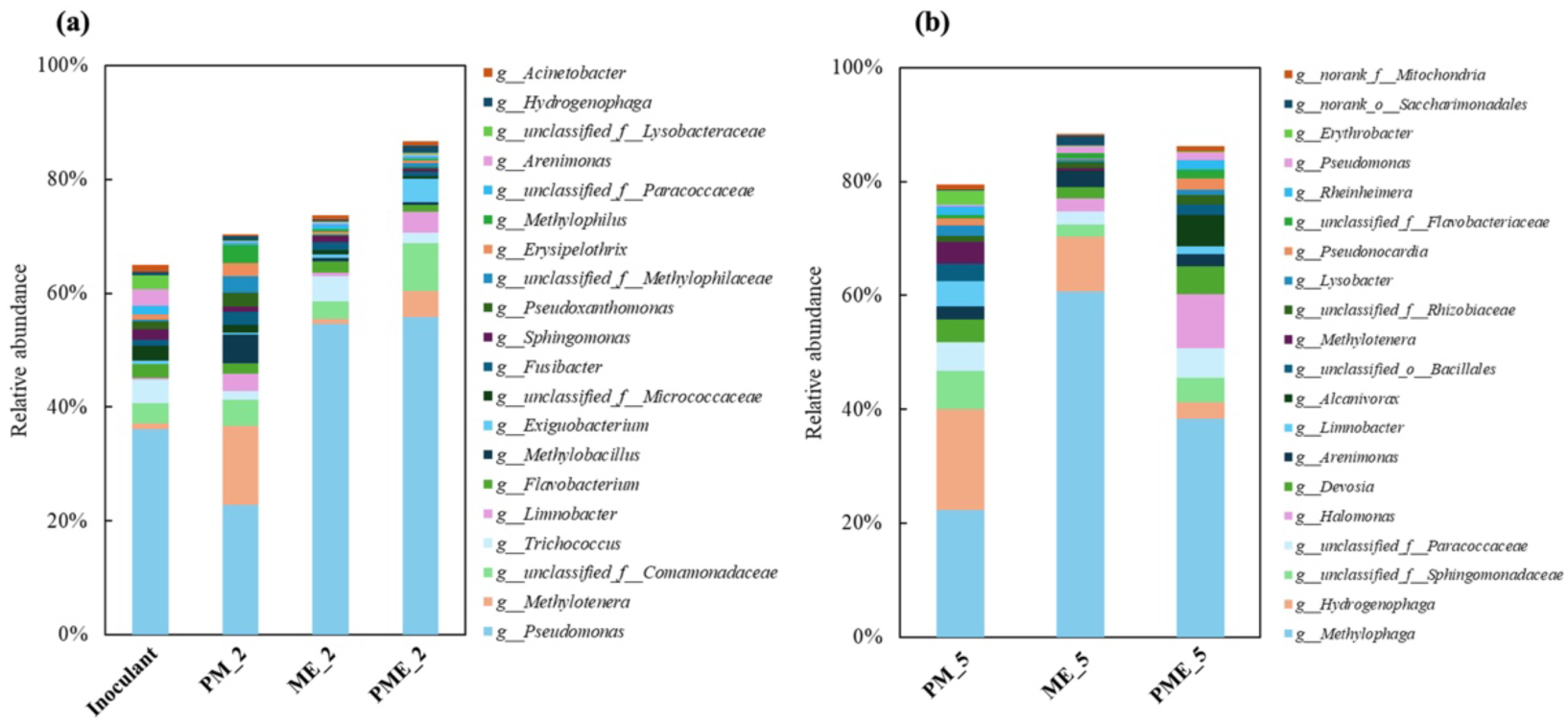
Microbial community composition in bulk samples from the column reactors. The 20 most abundant ASVs in (a) the inoculum and the PM, ME and PME treatments at the end of the batch experiment with an initial CCl_4_ concentration of 2 mg/L, and (b) the PM, ME and PME treatments at the end of the batch experiment with an initial CCl_4_ concentration of 5 mg/L. ASVs exhibiting significant differences in abundance relative to PM, identified using Fisher’s exact test, in (c) ME at 2 mg/L, (d) PME at 2 mg/L, (e) ME at 5 mg/L and (f) PME at 5 mg/L initial CCl_4_. PM, plants with live microbial inoculum; ME, live microbial inoculum under an applied electric field; PME, plants with live microbial inoculum under an applied electric field.

**Figure 5.**
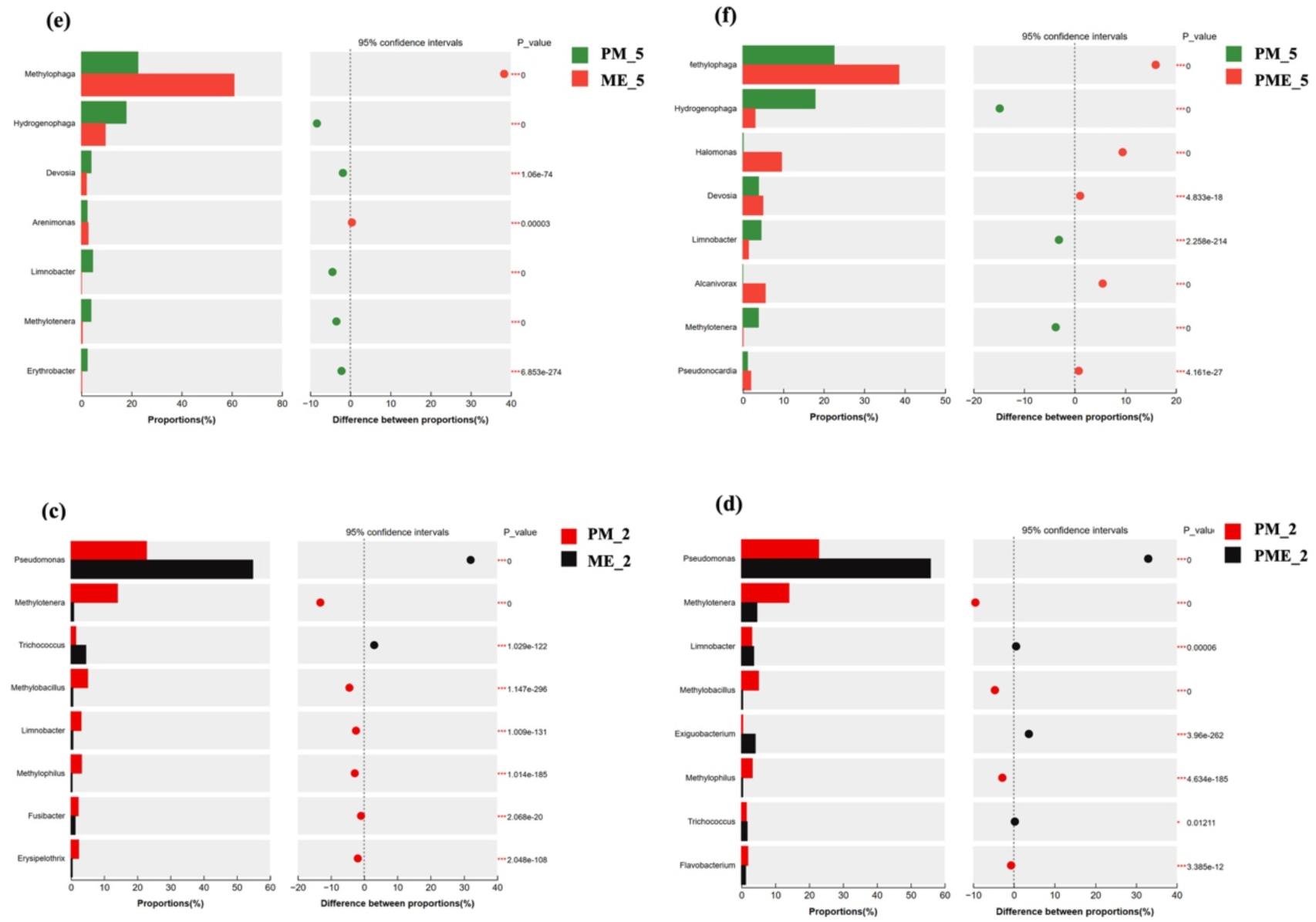
ASVs exhibiting significant differences in abundance relative to bulk samples, identified using Fisher’s exact test, in (a) ME anode at 2 mg/L, (b) ME cathode at 2 mg/L, (c) PME anode at 2 mg/L and (d) PME cathode at 2 mg/L initial CCl_4_.

**Figure 6.**
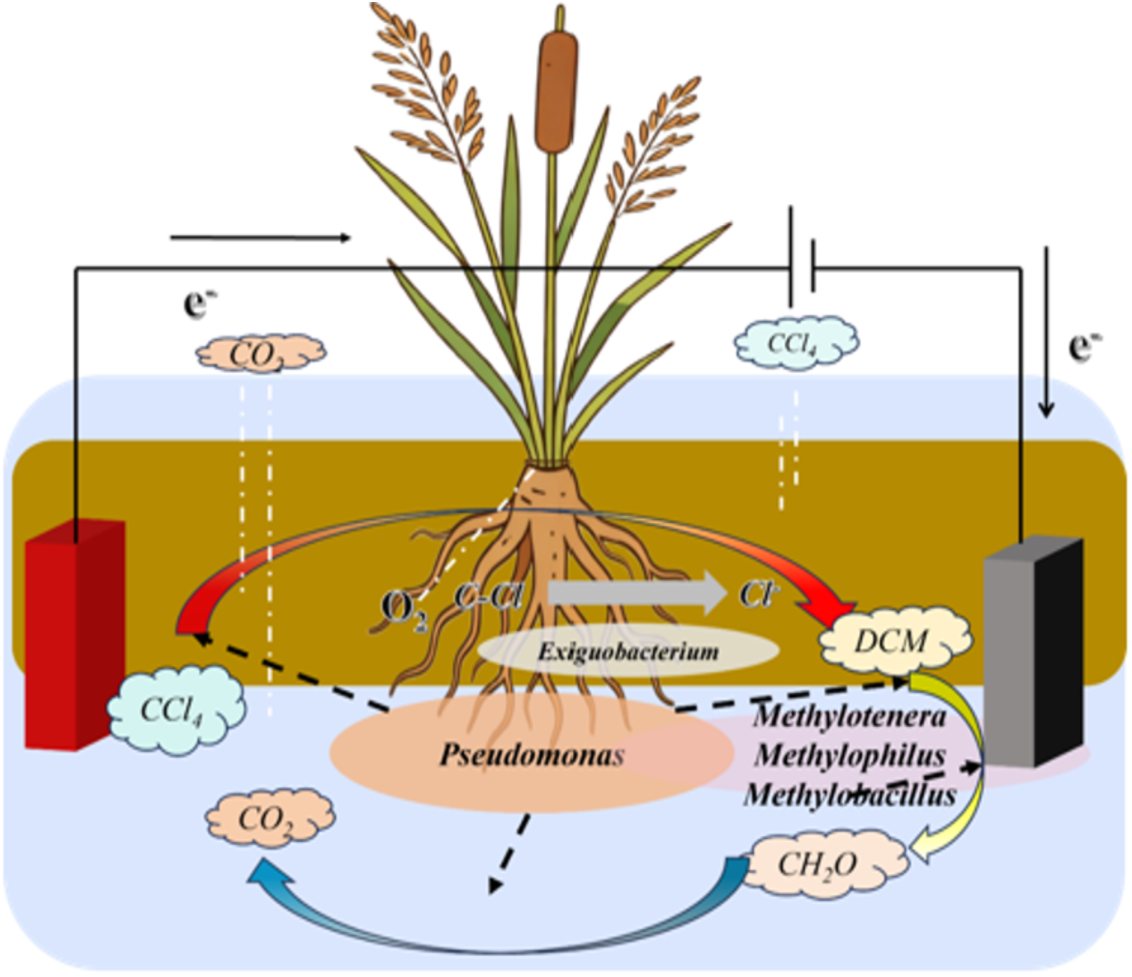
Proposed mechanisms of CCl_4_ removal in the plant–microbe system under an applied electric field.

The occurrence of multiple methylotrophic genera in bulk sediment and electrode-associated communities also suggests a potential role for C1-carbon metabolism. Methanol metabolism and associated carbon-assimilation pathways have been experimentally characterized in *Methylotenera mobilis* and a naturally occurring *Methylophaga*-like bacterium (Beck et al., 2011; Grob et al., 2015). These findings support considering methylotrophs as potential consumers of available C1 substrates. However, methanol introduced with the CCl_4_ stock solution provides an alternative explanation for their abundance. Their participation in the consumption of CCl_4_-derived intermediates therefore remains unresolved, and carbon assimilation should not be equated with complete mineralization. A potential electron-transfer role for *Exiguobacterium* is supported by electrochemical evidence from an iron-corrosion study involving *E. acetylicum* GD (Guo et al., 2025). This evidence provides a precedent for electron-transfer-related activity in an individual strain, but does not establish electrode-mediated CCl_4_ reduction by the *Exiguobacterium* populations detected here. Figure 6 therefore presents a proposed model connecting electrical stimulation, plant-associated conditions and potential microbial transformations. The bulk-sediment and electrode-community profiles identify candidate populations for further investigation, while their specific metabolic roles require functional validation. Shotgun metagenomics and gene-expression analyses of electrode-associated populations, combined with electrochemical measurements, transformation-product analysis and compound-specific isotope tracing, would help resolve the pathways and organisms responsible for CCl_4_ removal.

## 4. Discussion

### 4.1. Potential roles of electrical stimulation and plant–microorganism interactions

The results support investigating electrical stimulation and plant–microorganism interactions as complementary approaches to CCl_4_ removal. High removal in ME and PME across repeated cycles, together with treatment-associated ORP differences, suggests that electrical stimulation influenced conditions relevant to contaminant transformation. Cathodically generated hydrogen can support chloroform dechlorination, while complementary organisms ferment dichloromethane to acetate, demonstrating that subsequent transformation can also occur anaerobically (Fernández-Verdejo et al., 2022a). However, the electron-delivery pathway operating at the applied cell voltage of 0.6 V remains unresolved. Reeds may contribute by modifying microbial habitats and oxygen availability near roots. Spatially variable radial oxygen loss has been measured in Phragmites australis, providing a basis for considering localized oxygenation within otherwise oxygen-depleted sediments (Li et al., 2021). Such heterogeneity could support complementary reductive and subsequent oxidative transformations, depending on the intermediates and organisms present. The distinct bulk-sediment and electrode-associated communities identify candidate populations for testing these interactions. The coupling dechlorination with oxidation therefore remains a proposed mechanism requiring direct measurements of electrode potential, hydrogen, rhizosphere oxygen and transformation products.

### 4.2. Environmental implications and priorities for field application

Integrating electrical stimulation with planted sediment systems offers a potential treatment option for CCl_4_-contaminated groundwater at shallow, root-accessible interfaces. Sustained removal over repeated cycles supports further evaluation, while field applicability will depend on hydraulic contact, contaminant loading and seasonal plant performance. Field-scale reedbeds have treated groundwater containing 1,2-dichloroethane, demonstrating the practical relevance of planted treatment systems without establishing equivalent performance for CCl_4_ (Rahim et al., 2026). Permeable bioelectrochemical reactors provide another relevant precedent, although observed conversion of cis-dichloroethene to vinyl chloride highlights the importance of monitoring daughter products (Sassetto et al., 2024). Environmental benefit should therefore be assessed using contaminant mass balances, transformation-product toxicity and treatment durability, alongside removal efficiency. Further trials should use authentic groundwater and evaluate competing electron acceptors, co-contaminants, variable flow and extended operation. Measurements of electrical current, specific energy consumption and electrode maintenance requirements are also needed to establish operational feasibility. These assessments would determine whether the approach can deliver reliable contaminant attenuation without accumulating hazardous intermediates or imposing disproportionate resource demands.

## 5. Conclusion

This study demonstrated sustained CCl_4_ removal in sediment columns simulating wetland–groundwater interactions and identified microbial community responses to electrical stimulation and plant presence. At an initial concentration of 2 mg/L, PM, ME and PME achieved 79.5–82.9% removal within 48 h, exceeding AC and AP, which achieved 47.3% and 61.1%, respectively. Nearly complete removal occurred in ME and PME during the first cycle at 10 mg/L, with high removal maintained over subsequent cycles. Operation remained within a predominantly near-neutral to mildly alkaline pH range. Chloride increases accompanied CCl_4_ removal, whereas declining organic-carbon concentrations reflected changes in the broader carbon pool. Community profiling identified increased *Pseudomonas* abundance in PME and distinct electrode-associated populations, including *Methylobacillus* and *Methylophilus* in ME and *Methylotenera* in PME. These findings identify candidate organisms and support further development of electrically assisted planted systems. Validation through product and isotope analyses, electrochemical measurements and extended testing with authentic groundwater under variable flow and temperature is needed before field application.

## Acknowledgement

This study is supported by Open Fund (HNHSGE202506) of Engineering Research Center for Groundwater Resources and Environment of Hunan Province.

